# West Nile Virus Subverts T Cell Stimulatory Capacity of Human Dendritic Cells

**DOI:** 10.1101/602839

**Authors:** Matthew G. Zimmerman, James R. Bowen, Circe E. McDonald, Bali Pulendran, Mehul S. Suthar

## Abstract

West Nile virus (WNV) is a neurotropic flavivirus and the leading cause of mosquito-borne encephalitis in the United States. Recent studies in humans have found that dysfunctional T cell responses strongly correlate with development of severe WNV neuroinvasive disease. However, the contributions of human dendritic cells (DCs) in priming WNV-specific T cell immunity remains poorly understood. Here, we demonstrate that human monocyte-derived DCs (moDCs) support productive viral replication following infection with a pathogenic strain of WNV. Antiviral effector gene transcription was strongly induced during the log-phase viral growth, while secretion of type I interferons (IFN) occurred with delayed kinetics. Activation of RIG-I like receptor (RLR) or type I IFN signaling prior to log-phase viral growth significantly diminished viral replication, suggesting that activation of antiviral programs early can block WNV infection. In contrast to the induction of antiviral responses, WNV infection did not promote transcription or secretion of pro-inflammatory (IL-6, GM-CSF, CCL3, CCL5, CXCL9) or T cell modulatory cytokines (IL-4, IL-12, IL-15). There was also minimal induction of molecules associated with antigen presentation and T cell priming, including the co-stimulatory molecules CD80, CD86, and CD40. Functionally, WNV-infected moDCs dampened allogenic CD4 and CD8 T cell activation and proliferation. Combined, we propose a model where WNV subverts human DC activation to compromise priming of WNV-specific T cell immunity.

**Importance:** West Nile virus (WNV) is an encephalitic flavivirus that remains endemic in the United States. Previous studies have found dysfunctional T cell responses correlate to severe disease outcomes during human WNV infection. Here, we sought to better understand the ability of WNV to program human dendritic cells (DCs) to prime WNV-specific T cell responses. While productive infection of monocyte-derived DCs activated antiviral and type I interferon responses, molecules associated with inflammation and programming of T cells were minimally induced. Functionally, WNV-infected DCs dampened T cell activation and proliferation during an allogeneic response. Combined, our data supports a model where WNV infection of human DCs compromises WNV-specific T cell immunity.

## Introduction

West Nile virus (WNV) is a neurotropic flavivirus that remains the leading cause of mosquito-borne encephalitis in the United States (1). It is estimated that upwards of 6 million people have been infected by WNV in the US since its introduction in 1999, leading to over a thousand cases of neuroinvasive disease and nearly a hundred deaths each year (2). Following the bite of an infected mosquito, approximately 20% of individuals present with clinical outcomes ranging from mild febrile illness to severe neuroinvasive disease. Neuroinvasion is a serious complication with long term sequelae that includes ocular involvement, cognitive impairment, muscle weakness, and flaccid paralysis (3). The continued public health threat and lack of FDA-approved vaccines or specific therapeutics against WNV underpins the need to better understand the mechanisms of protective immunity during human infection.

The pathogenesis of human WNV infection is incompletely understood, although excellent mouse models have illuminated mechanisms of virus-induced encephalitis and critical features of immune control (4). The bite of an infected mosquito delivers high doses of WNV into the skin where keratinocytes, Langerhans cells, and dermal dendritic cells (DCs) are believed to be initial target cells of infection (5, 6). Over the next 24 hours, WNV migrates to the skin draining lymph nodes and replicates within resident DCs. Subsequent viremia promotes peripheral seeding of virus into permissive tissues such as the spleen, where DCs are targeted for infection (7). WNV then crosses the blood brain barrier and infects neurons within the central nervous system (CNS), leading to viral encephalitis. Restriction of viral replication by DCs during the early, peripheral phases of viral replication has been shown to be critical for limiting neuroinvasion and mitigating viral encephalitis (7, 8).

Within murine DCs, detection of WNV occurs primarily through the concerted efforts of RIG-I and MDA5 (9, 10), members of the retinoic acid inducible gene I (RIG-I) like receptor (RLR) family of cytosolic pattern recognition receptors. Signal transduction through the adaptor protein mitochondrial antiviral signaling (MAVS) triggers the nuclear factor-κB (NFκB) and interferon regulatory factor (IRF)-3, 5, and 7 dependent induction of type I interferon (IFN) and antiviral effector gene transcription (8). Following the MAVS-dependent secretion of type I IFN (10), signaling through the type I IFN receptor on DCs is required for early virus restriction and host survival (7).

In addition to direct restriction of viral replication, DCs are critical for the programming of antiviral CD8+ T cell responses that are required for clearance of WNV from the peripheral tissues and CNS (11). In humans, analysis of CD4+ and CD8+ T cells from the blood of WNV-infected patients has found dysfunctional T cell responses correlate with symptomatic disease outcome (12, 13). Decreased frequencies of CD4+ regulatory T cells also correlates with symptomatic WNV infection, highlighting the importance of a balanced T cell response (14). However, the contributions of human DCs in programming T cell immunity during human WNV infection remains poorly understood.

Here, we utilized primary human cells to demonstrate that WNV productively replicates within monocyte-derived DCs. Log-phase viral replication corresponded with induction of type I IFN and antiviral effector genes, with more delayed secretion of IFNα and IFNβ proteins. Activation of RLR or type I IFN signaling restricted viral replication, with RLR signaling remaining effective even after blockade of signaling through the type I IFN receptor. In contrast, WNV infection failed to up-regulate molecules involved in promoting inflammatory responses and priming of T cell immunity. Functionally, impaired DC activation resulted in diminished T cell proliferation by WNV-infected moDCs during an allogeneic response. Combined, WNV infection of human DCs activated antiviral responses while failing to program DCs to effectively prime WNV-specific T cell immunity.

## Results

### WNV productively infects human DCs

While DCs are an important cell type during infection with multiple flaviviruses, their contributions during human WNV infection remains limited. To model viral replication in human DCs, monocyte-derived DCs (moDCs) were generated from peripheral blood CD14+ monocytes and infected with a pathogenic strain of WNV (15). Viral replication was first detected at 12hpi, as noted by increased viral RNA synthesis **(Fig. 1A)**. Viral RNA levels continued to increase exponentially over the next 36hrs. Consistent with genome replication kinetics, release of infectious virus increased exponentially between 12 and 24hpi and plateaued at 48hpi **(Fig. 1B)**. Next, infected populations of moDCs were stained for intracellular expression of a structural protein found within the virus envelope (viral E protein) (16). Corresponding with log phase viral growth, the percentage of infected cells increased exponentially between 12 and 24hpi **(Fig. 1C)**. Infection plateaued between 24 and 48hpi, reaching upwards of 50% of cells positive for viral E protein. E protein expression was not observed in mock or UV inactivated virus infection controls. ImageStream analysis revealed that WNV E protein was localized predominantly within the cytoplasm and did not co-localize with the cell surface marker CD11c or the nucleus **(Fig. 1D)**. Declining percent infection at 72hpi corresponded with significant loss of cell viability **(Fig. 1E)**. Combined, three complementary measures of viral replication (viral RNA, infectious virus release, and viral E protein staining) confirm that human moDCs are productively infected by WNV with log phase viral growth beginning between 12 and 24hpi.

**Fig 1.**
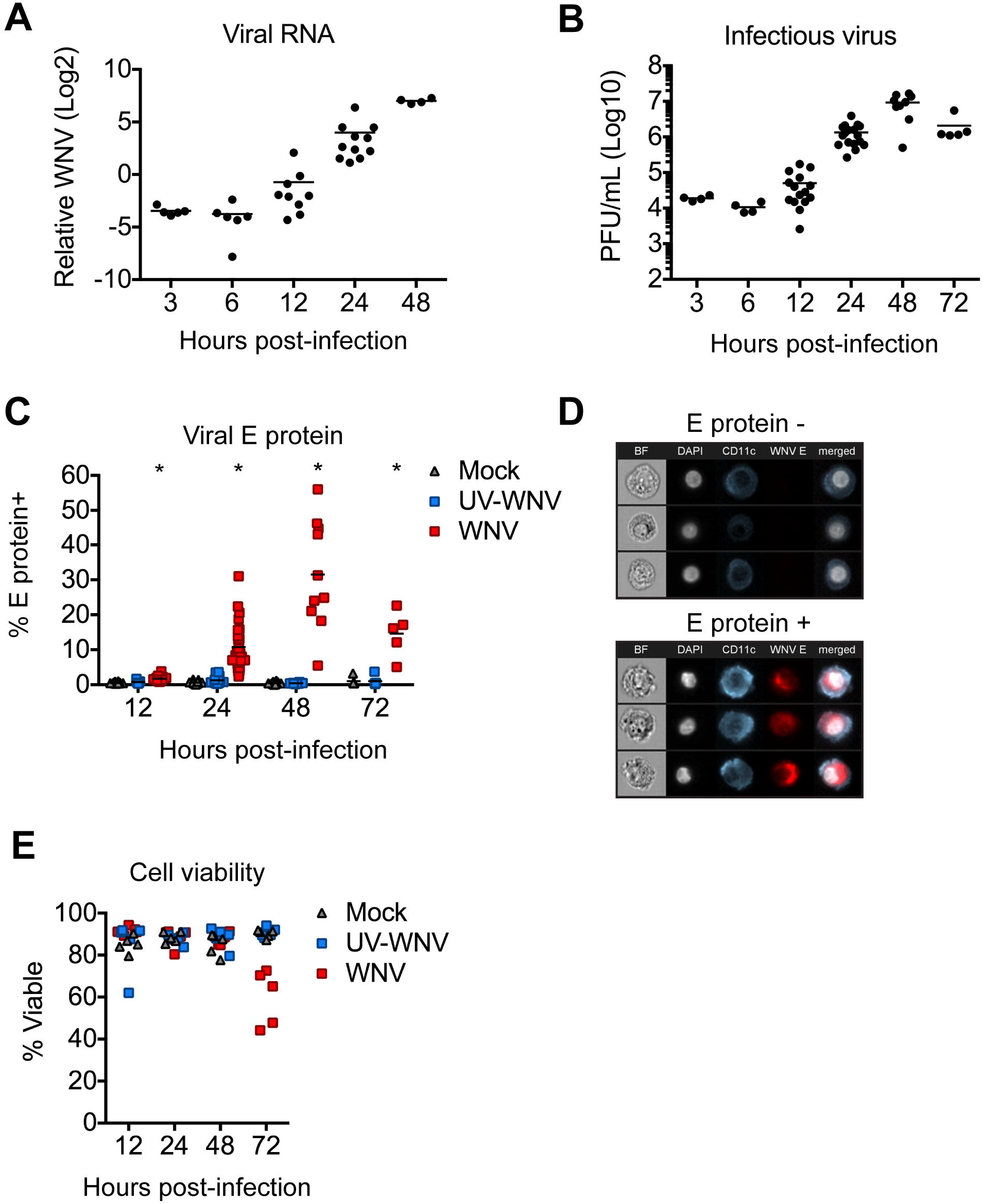
WNV productively infects human moDCs. moDCs were infected with WNV or UV-inactivated WNV (UV-WNV) at MOI 10 (as determined on Vero cells) and analyzed at indicated hours post-infection. **(A)** Viral RNA as quantitated in cell lysates by RT-qPCR. Shown as log_2_ normalized expression after normalization to *GAPDH*. Data is shown for each donor with the mean (n=5-11 donors). **(B)** Infectious virus release into the supernatant as determined by a viral plaque assay on Vero cells. Data is shown for each donor with the mean (n=4-17 donors). PFU, plaque-forming unit. **(C)** Percent E protein+ cells as determine by flow cytometry (Left panel). Data is shown for each donor with the mean (n=5-31 donors). **(D)** ImageStream analysis of WNV-infected moDCs labeled for viral E protein at 24hpi **(E)** Percent viable cells. Data is shown for each donor with the mean (n=5 donors).

### Innate immune signaling restricts WNV replication

Type I IFN within dendritic cells is critical for mediating protection against lethal infection outcome and controlling flavivirus replication (17, 18). We next determined the ability of both the RLR and type I IFN signaling pathways to restrict WNV infection of moDCs. We infected moDCs with WNV (MOI 10), treated cells with either a RIG-I agonist, MDA5 agonist, or IFNβ at 1 hpi and virus replication was measured at 24 hpi. We triggered RIG-I using a previously characterized and highly-specific agonist derived from the 3’ UTR of hepatitis C virus (19) and triggered MDA5 using high molecular weight poly(I:C), which preferentially activates MDA5 signaling upon delivery into the cytoplasm (20). Stimulation of RIG-I, MDA5, or IFN-β signaling potently restricted viral replication, with greater than 90% inhibition as measured by both viral burden in the supernatant and frequency of infected cells **(Fig. 2A)**. To confirm the role of type I IFN, we infected moDCs in the presence of an IFNAR2 blocking antibody and observed no effect on viral replication through 24 hpi, however, late viral control was compromised as shown by a 3-fold increase in the frequency of infected cells and a log-fold increase in viral replication at 48 hpi **(Fig. 2B)**. Notably, blocking type I IFN signaling in the presence of either RIG-I or MDA5 agonists still reduced viral replication, although we did observe a slight reduction in the efficiency of MDA5 signaling to reduce WNV infection in the presence of an anti-IFNAR2 neutralizing antibody **(Fig. 2C)** (21) Combined, these findings demonstrate that RIG-I, MDA5 and type I IFN signaling can efficiently block WNV replication in human DCs.

**Fig 2.**
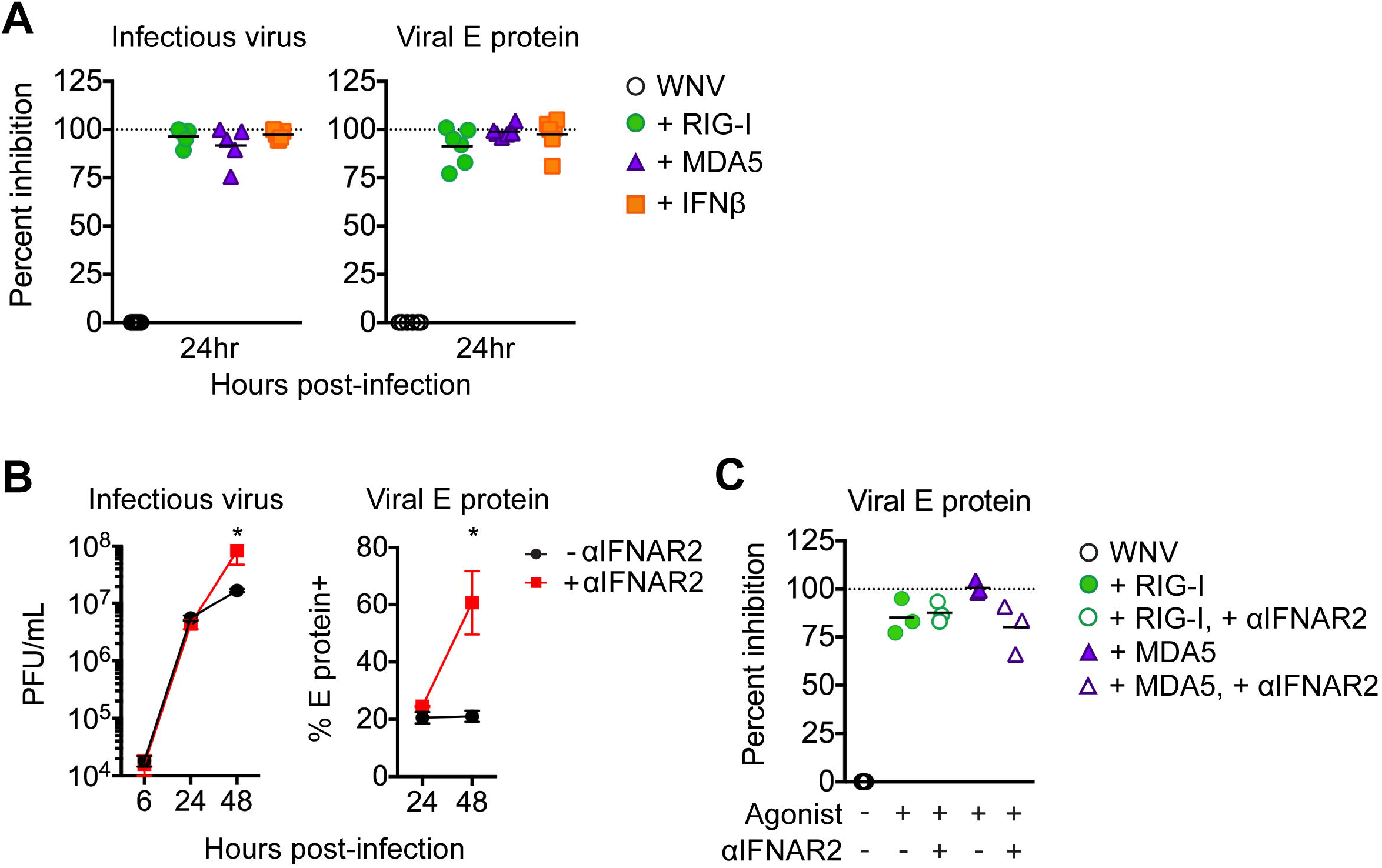
Innate immune signaling restricts WNV replication. **(A)** Experimental overview. moDCs were infected with WNV at MOI 10 (as determined on Vero cells) for 1hr and then treated with RIG-I agonist (100ng/1e6 cells), MDA5 agonist (100ng/1e6 cells), IFNβ (1000 IU/mL), or left untreated (“WNV”). **(B)** Infectious virus release into the supernatant (left panel) or viral E protein staining (right panel) was assessed at 24hpi. Data is represented as percent inhibition and shown for each donor with the mean (n= 5-6 donors). **(C)** moDCs treated as in B were incubated with or without anti-IFNAR2 (5µg/mL). For A and C, percent inhibition was calculated as: (1 - [WNV + agonist] / [WNV alone]) * 100. Dashed line indicates 100% inhibition, or complete block of viral infection. Data is shown for each donor with the mean (n= 3 donors). Statistical significance was determined as P < 0.05 using a Kruskal-Wallis test.

### WNV induces antiviral and type I IFN responses in human DCs

Traditional studies of antiviral responses have predominantly relied on approaches involving genetic ablation, gene knockdown, or gene overexpression methodologies (22). While useful, these approaches remain difficult to perform in primary human cells and may not accurately reflect the role of a given molecule during the normal course of infection. To overcome these limitations, we employed a systems biology approach to assess the antiviral landscape during WNV infection in human DCs **(Fig. 3A)**. We generated moDCs from 5 donors and performed messenger RNA sequencing following innate immune agonist treatment or infection with WNV. To study the early antiviral response during WNV infection, transcriptional responses were measured preceding (12 hpi) and during (24 hpi) log phase viral replication **(Fig. 1)**. Using weighted gene co-expression network analysis (WGCNA), we defined molecular signatures following stimulation of RIG-I, MDA5, or IFNβ signaling, identifying six clusters of co-expressed genes, or modules **(Fig. 3B)** (23). The module with the largest gene membership, module 5 (M5), was enriched for genes associated with the biologic process of “Defense response to virus”. Module 6 was also enriched for immune response related genes, while the remaining four modules were enriched for genes involved in biosynthetic processes, cellular metabolism, and stress responses. Given the large gene number and enrichment for antiviral response pathways, we focused our analyses on the M5 module.

**Fig 3.**
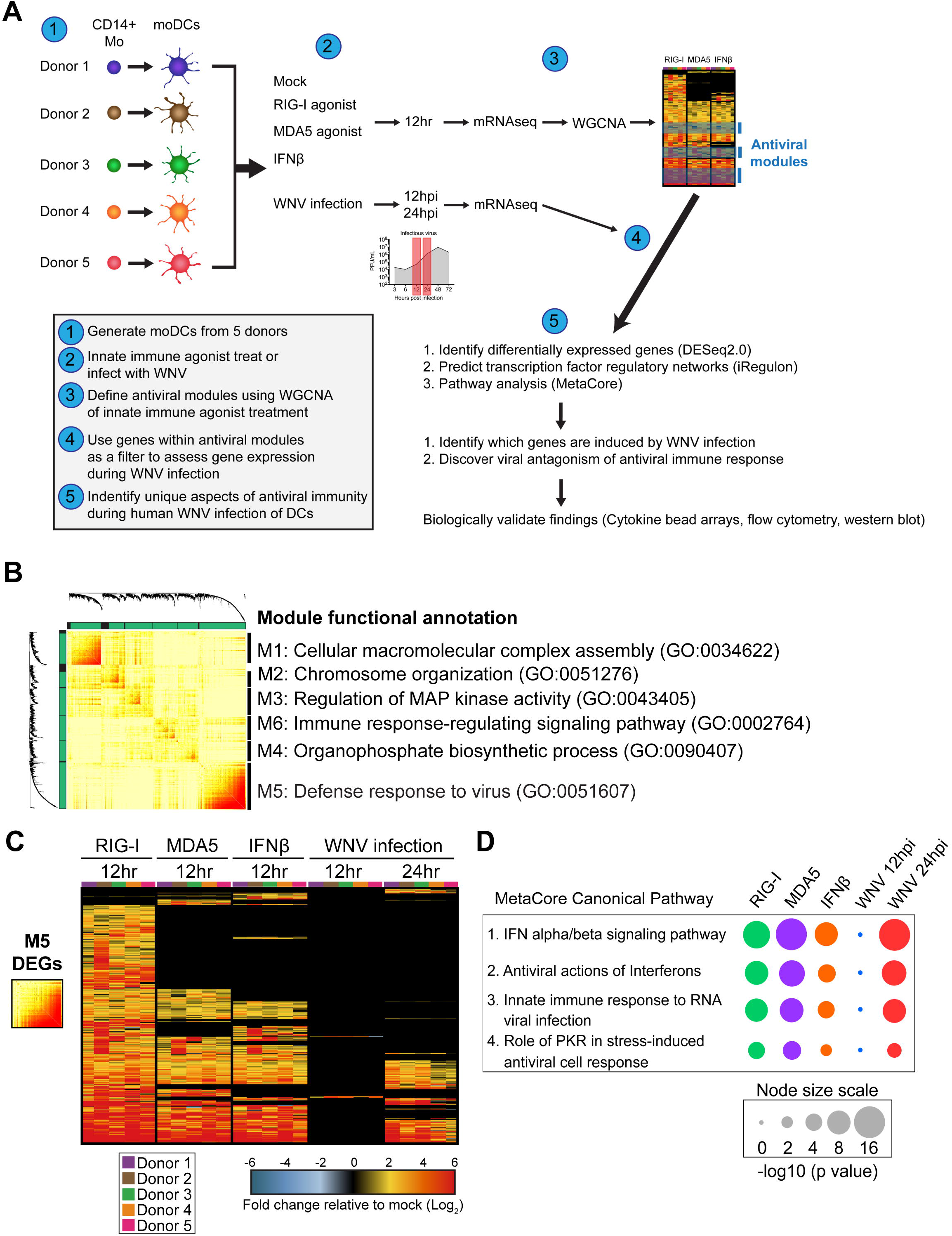
**(A)** Overview of systems biology approach used in this study. **(B)** Topologic overlap matrix showing enriched modules defined by WGCNA following 12 hr treatment with RIG-I agonist (100ng/1e6 cells), MDA5 agonist (100ng/1e6 cells), or IFNβ (1000 IU/mL). Functional annotation was performed using the DAVID Bioinformatics Resource version 6.8, with the top enriched biological process shown. **(C)** Heatmap of all module 5 differentially expressed genes with the log_2_ normalized fold change relative to uninfected, untreated cells shown. Genes that did not reach the significance threshold are depicted in black color. **(D)** Top enriched MetaCore canonical pathways of module 5 differentially expressed genes relative to uninfected and untreated cells (>2-fold change, p<0.01). Node size corresponds with the pathway enrichment significance score (–log_10_ p value) for each indicated treatment condition.

We next identified differentially expressed genes (DEGs) within the M5 module for each treatment condition, as compared to time-matched untreated and uninfected cells (>2-fold change, significance of p<0.01). RIG-I agonist treated cells induced a greater number of M5-related genes as compared to either MDA5 or IFNβ treated cells. In contrast, gene expression within the M5 module during WNV infection was temporally controlled: minimal gene expression at 12 hpi with more robust gene expression by 24 hpi **(Fig. 3C)**. MetaCore pathways enrichment analysis.of the M5 DEGs revealed four significantly enriched pathways including “IFN alpha/beta signaling”, “Antiviral activation of interferons”, “Innate immune response to RNA virus infection” and “Role of PKR in stress-induced antiviral cell response” (**Fig. 3D**). The expression patterns of host defense transcription factors, PRR signaling molecules, and antiviral effector genes were largely similar between the RIG-I agonist, poly(I:C), and IFNβ-treated DCs at 12hrs post stimulation, suggesting that upregulation of these genes are largely mediated through type I IFN signaling (**Fig. 4A**). Notably, WNV-infected DCs displayed minimal differentially expressed genes at 12hpi, however at 24hpi numerous antiviral effectors (e.g. *IFIT1, IFIT2, IFIT3, RSAD2, OASL*), molecules involved in RNA virus sensing (e.g. *DDX58*, *IFIH1*, *PKR*, *TLR3*), and the antiviral transcription factor *IRF7* were significantly up-regulated. Molecules involved in type I IFN signaling were also not induced at 12hpi but showed significant enrichment at 24hpi **(Fig. 4B)**. Despite enrichment of type I IFN genes at 24hpi, secretion of IFNα and IFNβ protein was not detected until 48hpi. The lack of detectable IFNα or IFNβ protein secretion until 48 hpi in human DCs is consistent with the significant increase in viral replication observed at 48 hpi when type I IFN signaling was blocked in WNV-infected DCs with an anti-IFNAR2 neutralizing antibody **(Fig. 2B)**. Combined, our data demonstrates that WNV infection of human DCs induces notable antiviral gene expression and that type I IFN signaling plays a role in late, but not early, restriction of viral replication

**Fig 4.**
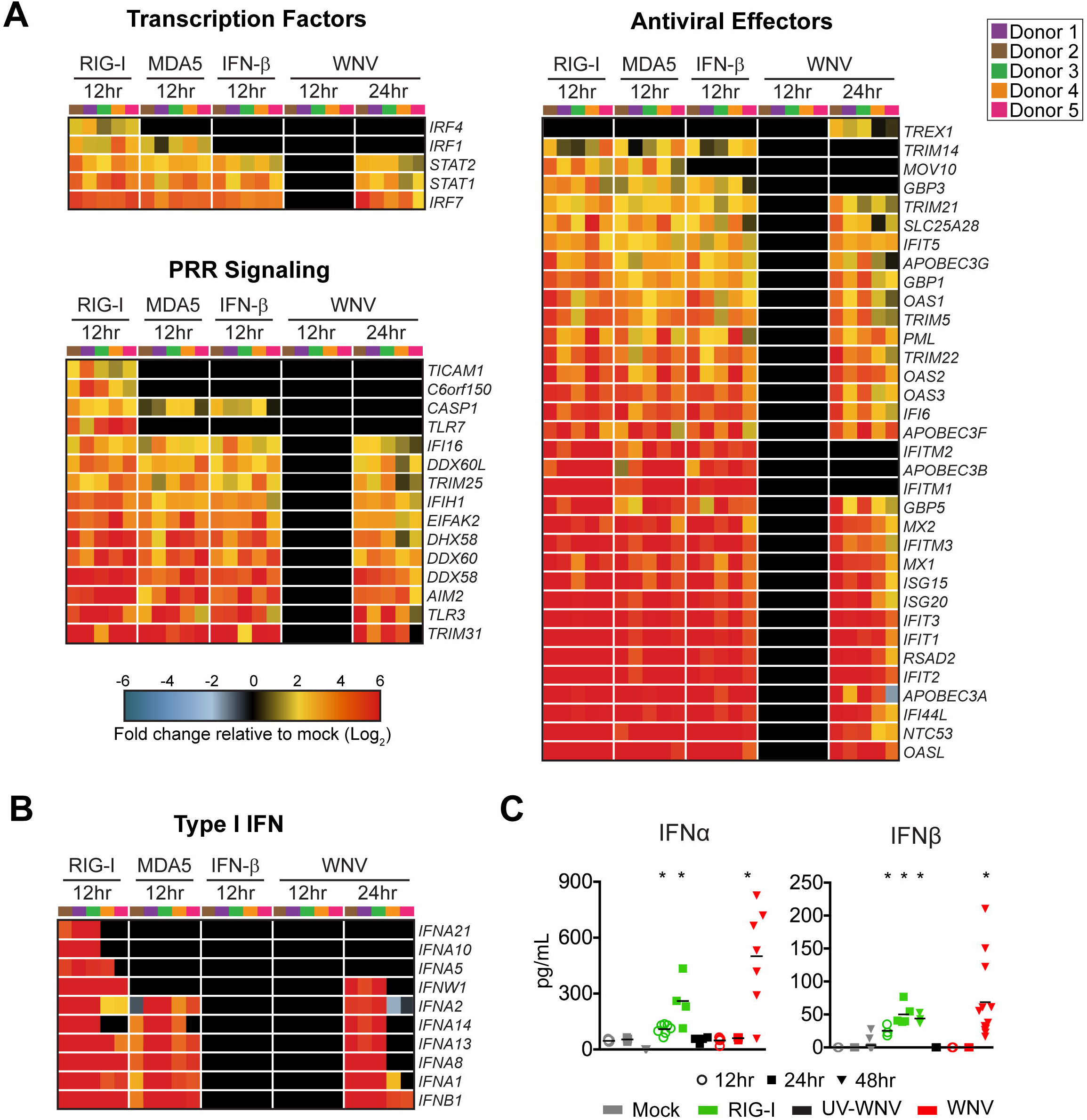
WNV induces robust antiviral and type I IFN responses. Messenger RNA sequencing was performed on moDCs generated from 5 donors after RIG-I agonist (100ng/1e6 cells for 12hrs), high MW poly(I:C) MDA5 agonist (100 ng/1e6 cells), or IFNβ (100 IU/mL) treatment or WNV infection (MOI 10; 12 and 24hpi). **(A)** Heatmap of differentially expressed genes (DEGs) corresponding to antiviral transcription factors, innate immune sensors, and antiviral effector genes. Genes that did not reach the significance threshold are depicted in black color. **(B)** Heatmap of DEGs corresponding to type I IFN responses. For all heatmaps, the log_2_ normalized fold change relative to uninfected, untreated cells is shown (>2-fold change, significance of p<0.01). Genes that did not reach the significance threshold are depicted in black color. Each column within a treatment condition is marked by a unique color and represents a different donor (n= 5 donors). **(C)** Secretion of IFNα and IFN β proteins into the supernatant following RIG-I agonist treatment (100ng/1e6 cells), infection with UV-inactivated WNV (MOI 10, “UV-WNV”), or infection with replication competent WNV (MOI 10, “WNV”). Data is shown for each donor with the mean (n= 4-11 donors). Statistical significance was determined as P < 0.05 using a Kruskal-Wallis test.

### WNV infection fails to promote inflammatory and T cell modulatory cytokine responses

We next assessed the induction of inflammatory cytokine and chemokine responses, an important component of antiviral immunity, DC activation, immune cell recruitment, and T cell priming (24, 25). In contrast to type I IFN and antiviral effector responses, WNV infection promoted minimal transcriptional enrichment of multiple cytokines involved in inflammatory cytokine responses (e.g. IL-15, IL-7, and IL-27), and chemotaxis (e.g. CCL2, CCL3, CCL4, CCL5, and CXCL9) **(Fig. 5A)**. *CXCR1* transcription was also selectively down-regulated during WNV infection. Importantly, RIG-I agonist treatment induced transcriptional expression of multiple inflammatory and T cell modulatory cytokines, confirming the ability of moDCs to mount pro-inflammatory responses upon innate immune stimulation. While RIG-I agonist induced inflammatory cytokines (IL-6 and GM-CSF), T cell promoting cytokines (IL-4, IL-15 and IL-12) and chemokines (CCL3, CCL5 and CXCL9), WNV-infected moDCs displayed little to no induction at the protein level of these cytokines/chemokines at 24 hpi **(Fig. 5B)**. These findings strongly suggest that WNV blocks the induction of proinflammatory cytokines/chemokines during infection.

**Fig 5.**
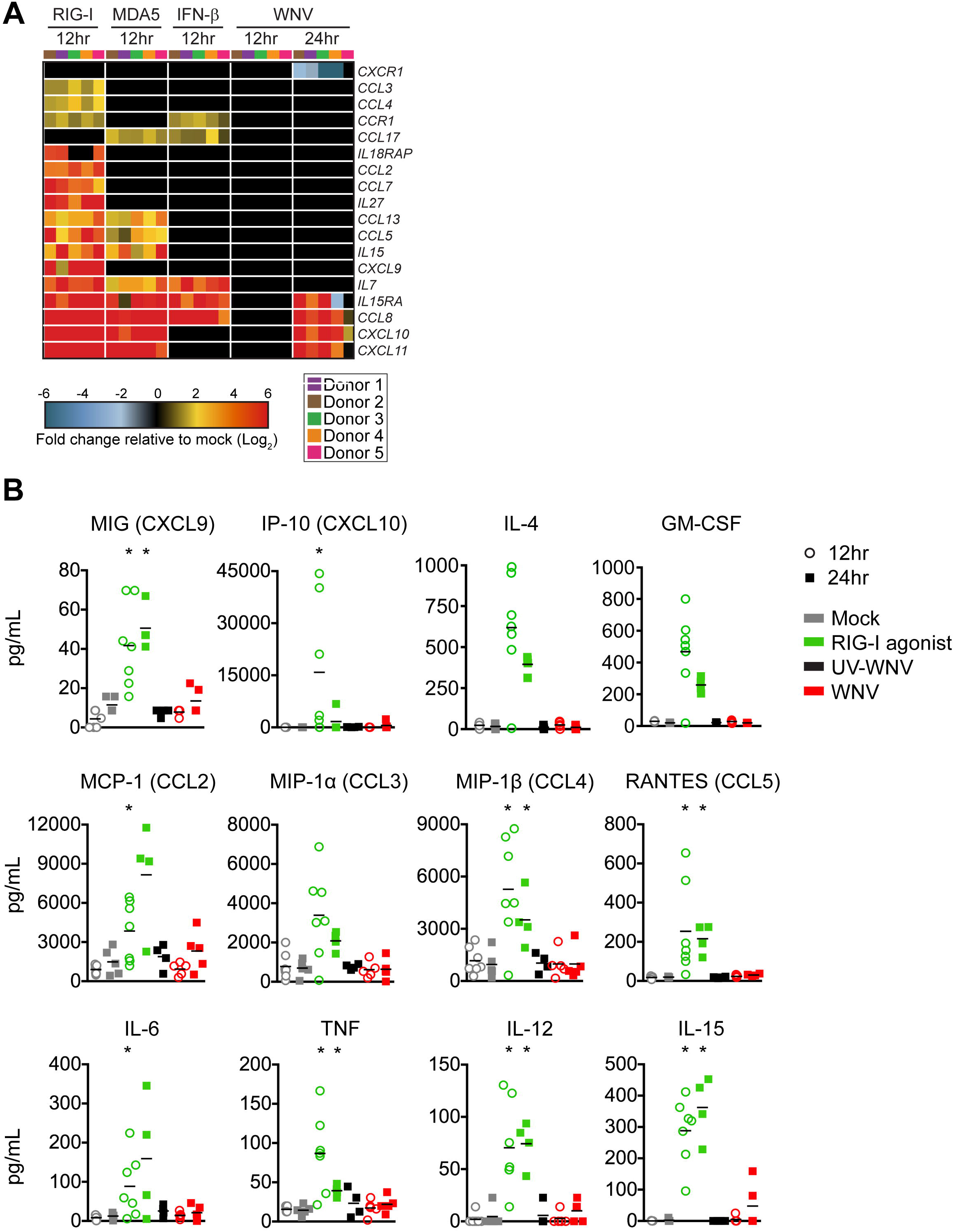
WNV infected DCs do not generate robust proinflammatory cytokine and chemokine responses. **(A)** Heatmap of genes involved in inflammatory cytokine responses and chemotaxis. The log_2_ normalized fold change relative to uninfected, untreated cells is shown (>2-fold change, significance of p<0.01). Genes that did not reach the significance threshold are depicted in black color. Each column within a treatment condition is marked by a unique color and represents a different donor (n= 5 donors). **(B)** Secretion of inflammatory cytokines, T cell modulatory cytokines, and chemokines were assessed by multiplex bead array following RIG-I agonist treatment (100ng/1e6 cells), infection with UV-inactivated WNV (MOI 10, “UV-WNV”), or infection with replication competent WNV (MOI 10, “WNV”). Responses were assessed at 24hr following treatment or infection. Data for each donor is shown with the mean (n=4-7 donors). Statistical significance was determined as P < 0.05 using a Kruskal-Wallis test.

### WNV infection does not induce molecules involved in T cell priming

In addition to the secretion of cytokines that modulate T cell responses, engagement of viral associated molecular patterns increases the surface expression of T cell co-signaling and MHC molecules on activated DCs (26). At the transcriptional level, WNV infection failed to induce multiple molecules associated with antigen presentation on MHC (*HLA-A, ERAP1*), *LAMP3* (27), proteasome subunits (*PSME1, PSMA2, PSMA4, PSMB10*), and *CD1D* (28) **(Fig. 6A)**. WNV also failed to significantly up-regulate genes involved in T cell co-signaling (e.g. *CD80, CD86, CD40*) and selectively up-regulated expression of galectin-9 (*LGALS9*), a ligand for the T cell inhibitory receptor TIM3 (12). These findings were biologically validated by flow cytometry, where WNV infection did not up-regulate cell surface levels of CD80, CD86, CD40 or MHC class II proteins within E protein+ cells at 24hpi or 48hpi **(Fig. 6B)**. Notably, high levels of WNV infection (MOI 100) still failed to induce expression of costimulatory and MHC class II molecules (**Fig. 6C**). In contrast to WNV infection, RIG-I agonist significantly up-regulated transcription of multiple molecules involved in antigen presentation and T cell co-signaling, corresponding with increased cell surface expression of CD80, CD86, CD40, and MHC II proteins. Combined, while WNV infection induces type I IFN and antiviral effector responses, WNV-infected DCs are compromised in their ability to induce inflammatory and chemotactic mediators important for immune activation, as well as antigen presentation and co-stimulatory molecules required for optimal T cell priming.

**Fig 6.**
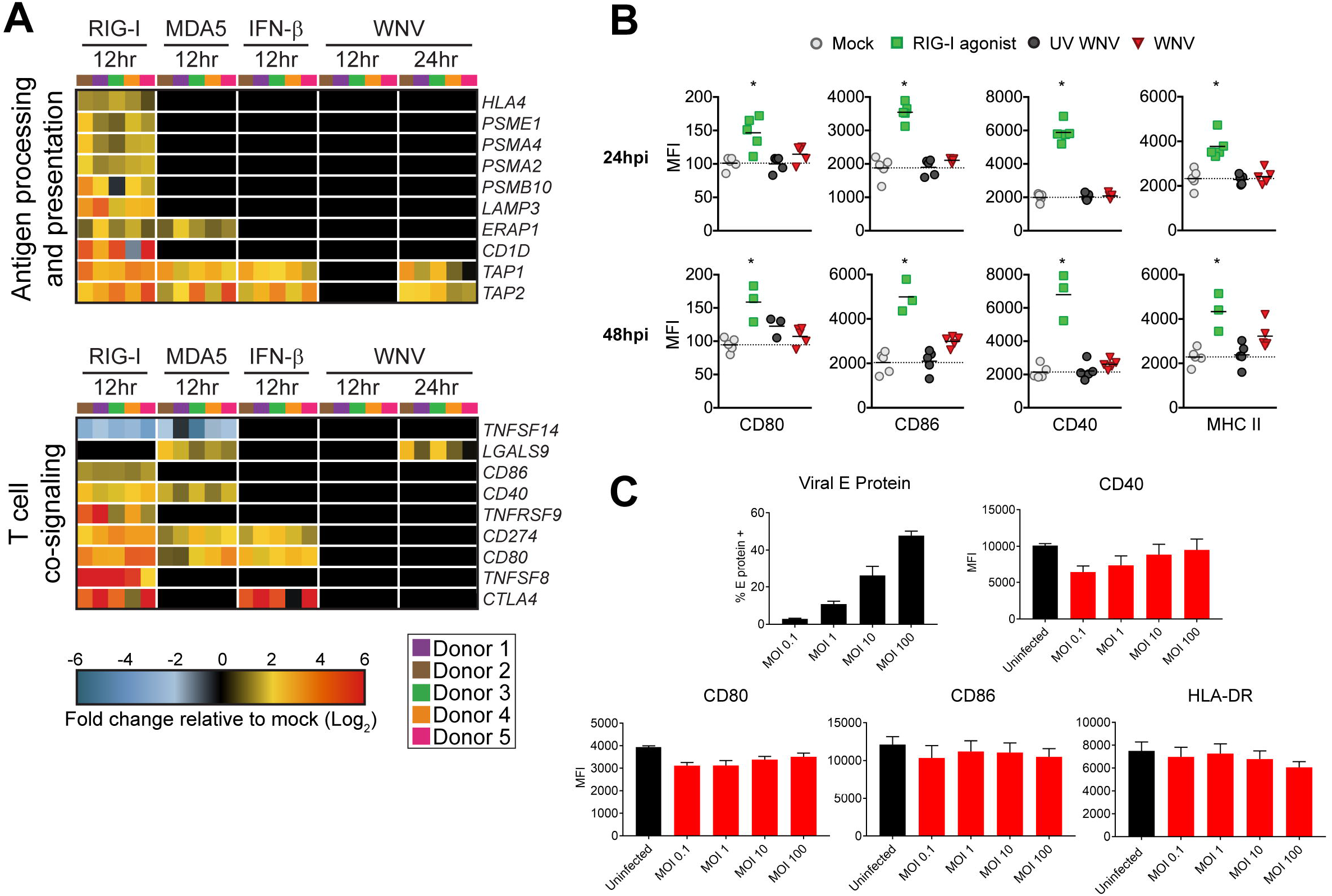
WNV-infected DCs fail to increase expression of molecules involved in antigen presentation and T cell co-stimulation. **(A)** Heatmap of genes involved in antigen processing and presentation or T cell co-signaling. The log_2_ normalized fold change relative to uninfected, untreated cells is shown (>2-fold change, significance of p<0.01). Genes that did not reach the significance threshold are depicted in black color. Each column within a treatment condition is marked by a unique color and represents a different donor (n= 5 donors). **(B)** Cell surface expression of CD80, CD86, CD40, or MHC II was quantitated by flow cytometry following RIG-I agonist treatment (100ng/1e6 cells), infection with UV-inactivated WNV (MOI 10, “UV-WNV”), or infection with replication competent WNV (MOI 10, “WNV”). Responses were assessed at 24hr and 48hr following treatment or infection. **(C)** Cell surface expression of CD80, CD86, CD40, or MHC II was quantitated by flow cytometry following infection with increasing MOIs of WNV at 24 hpi (MOI 0.1, 1, 10, and 100). For (B) and (C), WNV-infected moDCs were labeled for viral E protein and data is shown for the E protein+ population. Data for each donor is shown as median fluorescence intensity (MFI) with the mean (n=3-5 donors). Statistical significance was determined as P < 0.05 using a Kruskal-Wallis test.

### WNV-infected DCs dampen allogenic T cell proliferation

To determine if the minimal DC activation induced during WNV infection impairs T cell proliferation, we assessed the capacity of WNV-infected moDCs to drive an allogeneic T cell response. Uninfected moDCs induced notable activation of donor mismatched CD4 and CD8 T cells in a DC:T cell ratio dependent manner, as indicated by increased expression of the human T cell activation markers CD38 and HLA-DR (29) **(Fig. 7A)**. Allogenic activation of T cells corresponded with proliferation of upwards of 40% of CD4 and CD8 T cells in a DC:T cell ratio dependent manner **(Fig. 7B)**. In contrast, WNV infected moDCs diminished allogeneic CD4 and CD8 T cell activation, corresponding with significantly lower percentages of CD38+ HLA-DR+ and proliferated T cells. Combined, our data suggests that WNV infection induces minimal enrichment of molecules involved in DC activation, resulting in impaired T cell proliferation.

**Fig 7.**
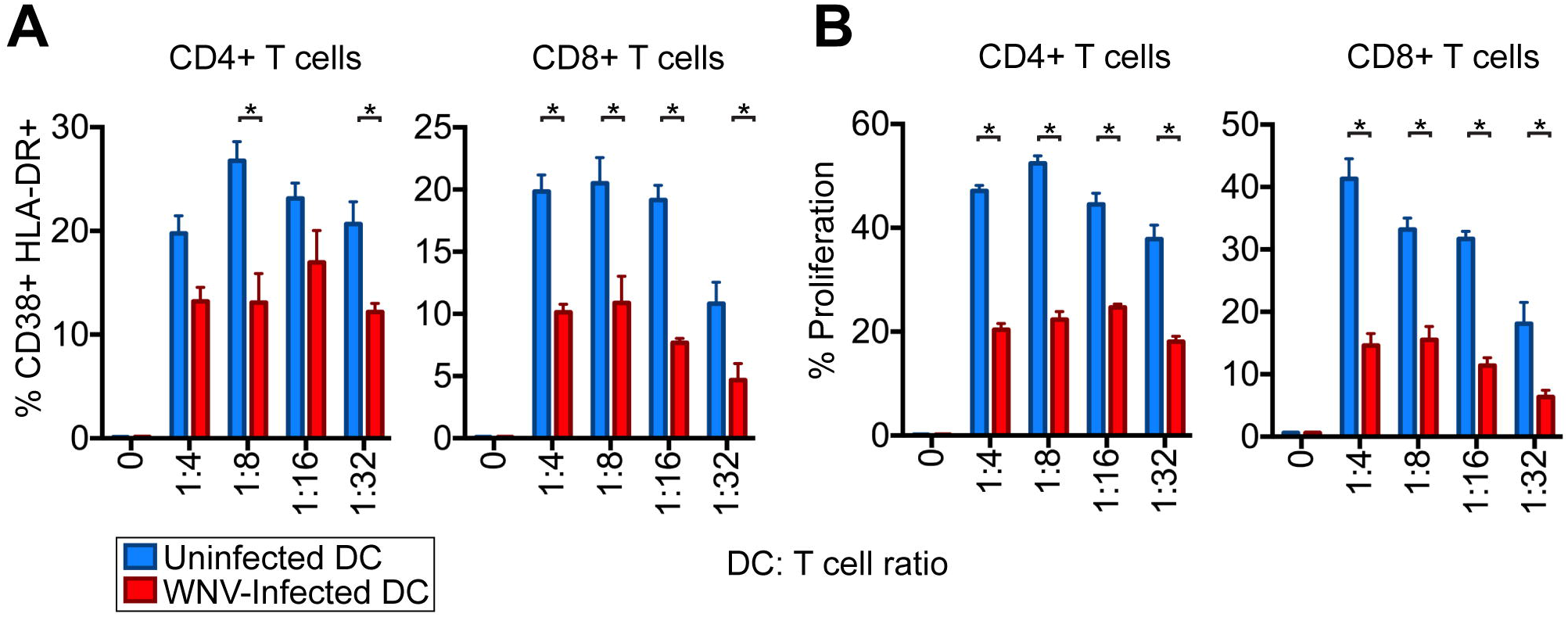
WNV-infected DCs are compromised in T cell proliferation. moDCs were left uninfected or infected with WNV (MOI 10) for 24hrs. Allogeneic CD4 or CD8 T cells were labeled with CellTrace violet (CTV) and incubated with uninfected or WNV infected moDCs at the indicated DC:T cell ratios in the presence of an anti-E16 WNV blocking antibody to limit spreading infection (5µg/mL) for 6 days. **(A)** The percentage of cells double positive for the T cell activation markers CD38 and HLA-DR on day 6 of allogeneic co-culture. **(B)** The percentage of cells that had proliferated by day 6 of allogeneic co-culture. Percent proliferation was defined as any cell that diluted CTV as compared to a “no DC, T cell only control”. Statistical significance was determined as P < 0.05 using a two-way ANOVA analysis.

## Discussion

In this study, we combined virologic and immunologic measures with transcriptomic analysis to better understand antiviral responses during WNV infection in primary human DCs. WNV productively infected human moDCs and induced cell death, coinciding with declining viral growth kinetics. RIG-I, MDA5, and IFNβ signaling potently restricted viral replication, corresponding with strong activation of antiviral defense response genes. In contrast, there was minimal up-regulation of inflammatory mediators or molecules involved in T cell priming. Functionally, WNV-infected moDCs promoted impaired allogeneic T cell proliferation and activation.

Studies in mice have found that RLR and type I IFN signaling are critical for viral restriction and host survival during WNV infection, however, the contributions of innate immune signaling during infection of human cells remains limited (7, 30). Here, we demonstrated that RIG-I, MDA5, and IFNβ signaling potently restrict WNV replication through induction of strong antiviral gene transcription, suggesting that similar to mice, RLR and type I IFN signaling are important for viral control during human WNV infection. RIG-I and MDA5 agonists also remained efficient in blocking WNV replication independent of type I IFN signaling, consistent with the ability of RLR signaling to induce antiviral gene expression in the absence of the type I IFN receptor in mice (30). Combined, our results confirm the importance of RLR and type I IFN signaling in the induction of antiviral responses and restriction of viral replication within primary human DCs.

These findings are similar to previous work, where WNV infection also failed to induce inflammatory cytokine secretion (31). Infection of moDCs with a non-pathogenic WNV isolate, WNV Kunjin, also induced minimal production of IL-12, despite notable up-regulation of both CD86 and CD40 (28). This suggests that an inability to induce inflammatory cytokine responses may be shared among WNV strains, while pathogenic strains have evolved unique mechanisms to subvert antigen presentation and T cell activation. The failure of WNV to activate human moDCs is also similar to our recent work with ZIKV (32). In contrast to WNV and ZIKV, infection of moDCs with the yellow fever virus vaccine strain (YFV-17D) up-regulates multiple inflammatory mediators and surface expression of CD80 and CD86 (26). The ability of YFV-17D to induce strong DC activation may reflect the loss of a viral antagonist during the attenuation process, similar to the ability of WNV Kunjin to induce up-regulation of CD86 and CD40 (28). Alternatively, the ability of YFV-17D to induce DC activation may be an inherent property of certain flaviviruses. Indeed, DENV has also been found to activate inflammatory responses and up-regulate co-stimulatory molecules following infection (33, 34). Altogether, our recent studies may suggest that certain neurotropic flaviviruses are particularly adept at subverting DC:T cell signaling.

Due to the largely subclinical presentation of WNV infection in humans, understanding genetic correlates of susceptibility and viral restriction remains exceedingly difficult. However, modeling WNV infection in mice lacks the genetic variation seen within outbred human populations. To combat this issue, the collaborative cross (CC) mouse model system, recombinant inbred mice containing genetic diversity from eight founder mouse strains, has been recently developed to study host antiviral responses within a genetically diverse population (35). Using the CC mouse system, one group observed increased regulatory T cells (Tregs) infiltration and no immunopathology in the brains of asymptomatic WNV-infected mice (36). This corroborated well to earlier human studies showing that increased levels of Tregs correlated to improved outcomes during WNV infection (14). These studies have focused largely on observing WNV-specific T and B cell responses to in the CC model system; however, the effects of these diverse polymorphisms on DC function during WNV infection remains largely untouched. Similar to our current study in human moDCs, transcriptomic analyses from whole spleens and brains of WNV-infected CC mice have also shown differences in antigen presentation, T cell signaling, and inflammatory cytokine production (37, 38). Altogether, the CC mouse system can be utilized in future studies to recapitulate human disease and understand DC responses during WNV infection.

An important observation of our study was that WNV infection did not trigger DC activation, as determined by upregulation of costimulatory protein expression. Through an allogeneic T cell assay, we found that WNV-infected moDCs were less efficient with inducing CD4+ and CD8+ T cell proliferation as compared to mock-infected moDCs. WNV-specific T cell responses have been detected in both symptomatic and asymptomatic WNV infection in humans (39). However, quality rather than quantity of the CD4+ and CD8+ T cell responses during WNV infection is an important predictor of symptomatic infection outcome. Dysregulated Th1 CD4+ T cell responses were found to strongly correlate with neuroinvasive disease (13). Additionally, decreased numbers of regulatory T cells have also been implicated in symptomatic and neuroinvasive infection in WNV-infected individuals, suggesting that immunomodulation of WNV-specific T cells responses are essential for avoiding immunopathology (14). Lastly, a recent study linked expression of the inhibitory T cell receptor Tim-3 on T cells with progression to symptomatic disease outcome (12). Combined, these findings demonstrate that development of an effective T cell response is critical for modulating infection outcome (symptomatic vs asymptomatic) during WNV infection. Our studies have now determined that WNV interferes with DC activation, through inhibition of costimulatory molecule expression and pro-inflammatory cytokine production, which can lead to dysregulated T cell responses, immunopathology and excessive neuronal injury.

In summary, our systems biology approach defined the antiviral landscape seen during RLR and type I IFN signaling as well as WNV infection in human DCs. Through our study, we observed that WNV can downregulate numerous genes responsible for establishing proper WNV-specific adaptive immune responses in human DCs, negatively affecting proper CD4+ and CD8+ T cell responses. Altogether, our study significantly advances our understanding of how WNV disrupts antiviral immunity during human infection.

## Materials and Methods

### Ethics statement

Human peripheral blood mononuclear cells (PBMCs) were obtained from de-identified healthy adult blood donors and processed immediately. All individuals who participated in this study provided informed consent in writing in accordance to the protocol approved by the Institutional Review Board of Emory University, IRB#00045821, entitled “Phlebotomy of healthy adults for the purpose of evaluation and validation of immune response assays”.

### Viruses

WNV stocks were generated from an infectious clone, WNV isolate TX 2002-HC, and passaged once in Vero cells, as previously described (15). WNV stocks were titrated on Vero cells by plaque assay. moDCs were infected with WNV at MOI 10 for 1hr at 37°C in cRPMI (without GM-CSF or IL-4). After 1hr, virus was washed off, cells were resuspended in fresh cRPMI, and incubated at 37°C for 3-72 hours.

### Cell lines

Vero cells (WHO Reference Cell Banks) were maintained in complete DMEM. Complete DMEM was prepared as follows: DMEM medium (Corning) supplemented with 10% fetal bovine serum (Optima, Atlanta Biologics), 2mM L-Glutamine (Corning), 1mM HEPES (Corning), 1mM sodium pyruvate (Corning), 1x MEM Non-essential Amino Acids (Corning), and 1x Antibiotics/Antimycotics (Corning). Complete RPMI was prepared as follows: cRPMI; RPMI 1640 medium (Corning) supplemented with 10% fetal bovine serum (Optima, Atlanta Biologics), 2mM L-Glutamine (Corning), 1mM Sodium Pyruvate (Corning), 1x MEM Non-essential Amino Acids (Corning), and 1x Antibiotics/Antimycotics (Corning).

### Generation of monocyte derived dendritic cells

To generate human moDCs, CD14+ monocytes were differentiated in cRPMI supplemented with 100ng/mL of GM-CSF and IL-4 for 5-6 days, as previously described (32). In brief, freshly isolated PBMCs obtained from healthy donor peripheral blood (lymphocyte separation media; StemCell Technologies) were subjected to CD14+ magnetic bead positive selection using the MojoSort Human CD14 Selection Kit (BioLegend). Purified CD14+ monocytes were cultured in complete RPMI supplemented with 100ng/mL each of recombinant human IL-4 and GM-CSF (PeproTech) at a cell density of 2e6 cells/mL. After 24hr of culture, media and non-adherent cells were removed and replaced with fresh media and cytokines. Suspension cells (“moDCs”) were harvested after 5-6 days of culture and were consistently CD14-, CD11c+, HLA-DR+, DC-SIGN+, and CD1a+ by flow cytometry. For experimentation, moDCs were maintained in complete RPMI without GM-CSF or IL-4.

### Quantitative reverse transcription-PCR (qRT-PCR)

Total RNA was purified (Quick-RNA MiniPrep Kit; Zymo Research) and viral RNA was reverse transcribed (High Capacity cDNA Kit; Applied Biosystems) using 1 pmol of a GVA tagged (underlined) primer (5’-<u>TTTGCTAGCTTTAGGACCTACTATATCTACCT</u>GGGTCAGCACGTTTGTCATTG-3’) directed against the E gene (18, 40). Reverse transcribed viral sequences were detected by qRT-PCR (TaqMan Gene Expression Master Mix; Applied Biosystems) using 10 pmol of primers (5’-TTTGCTAGCTTTAGGACCTACTATATCTACCT3’ and 5’-TCAGCGATCTCTCCACCAAAG-3’) and 2.5 pmol of hydrolysis probe (5’-FAM-TGCCCGACCATGGGAGAAGCTC-3IABkFQ-3’). All custom primers and probes were obtained from Integrated DNA Technologies. All qRT-PCR was normalized to the amount of GAPDH (Hs02758991_g1; Applied Biosystems) in each respective sample.

### Quantitation of infectious virus

Infectious virus was quantitated using a plaque assay on Vero cells with a 1% agarose overlay and crystal violet counterstain, as previously described (15).

### Innate immune agonists

To stimulate RIG-I signaling, 100ng of RIG-I agonist derived from the 3’-UTR of hepatitis C virus (19) was transfected per 1e6 cells using TransIT-mRNA transfection kit (Mirus). For stimulation of MDA5 signaling, 100ng of high molecular weight poly-(I:C) was transfected per 1e6 cells using LyoVec transfection reagent (Invivogen). To stimulate type I IFN signaling, cells were incubated with 100 IU/mL of human recombinant IFNβ. In select experiments, different doses of agonists were used and this is indicated within the respective figure legend. To inhibit type I IFN signaling, 5µg/mL anti-human Interferon-α/β Receptor Chain 2 (MMHAR-2; EMD Milipore) blocking monoclonal antibody was used.

### RNA sequencing and bioinformatics

moDCs were generated from 5 donors and either treated with innate immune agonists for 12hr (RIG-I, MDA5, or IFNβ) or infected with WNV (12hpi and 24hpi). Total RNA was purified (Quick-RNA MiniPrep Kit; Zymo Research) and mRNA sequencing libraries were prepared for RNA sequencing (Illumina TruSeq chemistry). RNA sequencing was performed on a Illumina HiSeq 2500 System (100bp single end reads). Sequencing reads were mapped to the human reference genome 38. Reads were normalized and differential expression analysis performed using DESeq2 (41). Differentially expressed genes were determined by 2-fold change and *P*<0.01. The raw data of all RNA sequencing will be deposited into the Gene Expression Omnibus (GEO) repository and the accession number will be available following acceptance of this manuscript. Weighted gene co-expression module analysis was performed on DESeq2 normalized mapped reads (TIBCO Spotfire with Integromics Version 7.0) from RIG-I agonist, MDA5 agonist, IFNβ, and mock treated samples. First, the datasets were reduced to focus the network analysis on the 5446 most variable genes (as determined by variation value greater than 1) using the Variance function in R. We constructed a signed weighted correlation network by generating a matrix pairwise correlation between all annotated gene pairs. The resulting biweight mid-correlation matrix was transformed into an adjacency matrix using the soft thresholding power (β1) of 12. The adjacency matrix was used to define the topological overlap matrix (TOM) based on a dissimilarity measurement of 1- TO. Genes were hierarchically clustered using average linkage and modules were assigned using the dynamic tree-cutting algorithm (module eigengenes were merged if the pairwise calculation was larger than 0.75). This resulted in the construction of six modules.

### Flow cytometry

Cells were prepared for analysis as previously described (32). In brief, cells were Fc receptor blocked for 10 min, stained for phenotypic and activation markers for 20 min, and viability stained for 20 min (Ghost Dye Violet 510, Tonbo Biosciences). For intracellular staining of WNV E protein, cells were fixed and permeabilized (Transcription Factor Staining Buffer Kit, Tonbo Biosciences) and labeled with E16-APC for 20min at room temperature (16). Flow cytometry data was analyzed using FlowJo version 10 software. ImageStream data was analyzed using the Amnis IDEAS software. For moDC studies, the following antibody clones from Biolegend were used: CD11c (B-Ly6), CD80 (2D10), CD86 (IT2.2), CD40 (5C3), HLA-DR (G46-6; BD Bioscience), CD14 (M5E2), CD1a (HI149).

### T cell proliferation assay

Freshly isolated PBMCs obtained from healthy donor peripheral blood (lymphocyte separation media; StemCell Technologies) were subjected to CD4 or CD8 T cell magnetic bead negative selection using the MojoSort Human CD4 or CD8 Selection Kit (BioLegend). Isolated CD4 or CD8 T cells were labeled with CellTrace Violet (CTV) Cell Proliferation Kit (ThermoFisher) per the manufacturer’s instructions. In a 96-well U bottom plate, CTV labeled CD4 or CD8 T cells (2e5 ells) were mixed with different ratios of either uninfected moDCs, or moDCs infected with WNV for 24hr (1:4, 1:8, 1:16, 1:32, 1:64, and 1:128 DC:T cell ratios). To prevent spreading infection, we added anti-E16 neutralizing antibody at 5µg/mL throughout the DC:T cell co-culture period (16). After 6 days of co-culture, cells were stained for surface expression of CD4 or CD8, CD3, CD38, and HLA-DR. Proliferation, by CTV dilution, and T cell activation (CD38+HLA-DR+) were assessed by flow cytometry (42).

### Multiplex bead array

Cytokine analysis was performed on supernatants using a human 25-plex panel (ThermoScientific) and a custom 2-plex panel with human IFNβ and IFNα simplex kits (eBioscience) as described previously (32). Cytokines analyzed included: included: IFN-α, IFNβ, GM-CSF, TNF-α, IL-4, IL-6, MIP-1α, IL-8, IL-15, IL-2R, IP-10, MIP-1β, Eotaxin, RANTES, MIG, IL-1RA, IL-12 (p40/p70) IL-13, IFN-γ, MCP-1, IL-7, IL-17, IL-10, IL-5, IL-2, and IL-1β.

### Statistics

All statistical analysis was performed using GraphPad Prism version 6 software. The number of donors varied by experiment and is indicated within the figure legends. Statistical significance was determined as P<0.05 using a Kruskal-Wallis test (when comparing more than two groups lacking paired measurements), a Wilcoxon test (when comparing two groups with paired measurements), or a two-way ANOVA (when comparing two groups across multiple independent variables). All comparisons were made between treatment or infection conditions with a time point matched, uninfected and untreated control.

## Acknowledgements

We thank the Children’s Healthcare of Atlanta and Emory University Pediatric Flow Cytometry Core for providing access to flow cytometry, ImageStream, and Luminex systems, and the Yerkes Genomics Core for performing RNA sequencing.

